# Hostile: accurate host decontamination of microbial sequences

**DOI:** 10.1101/2023.07.04.547735

**Authors:** Bede Constantinides, Martin Hunt, Derrick W Crook

## Abstract

**Motivation:** Microbial sequences generated from clinical samples are often contaminated with human host sequences that must be removed for ethical and legal reasons. Care must be taken to excise host sequences without inadvertently removing target microbial sequences to the detriment of downstream analyses such as variant calling and *de novo* assembly.

**Results:** To facilitate accurate host decontamination of both short and long sequencing reads, we developed Hostile, a tool capable of accurate host read removal using a laptop. We demonstrate that our approach removes at least 99.6% of real human reads and retains at least 99.989% of simulated bacterial reads. Using Hostile with a masked reference genome further increases bacterial read retention (>=99.997%) with negligible (<=0.001%) reduction in human read removal performance. Compared with an existing tool, Hostile removes 21-23% more human short reads and 22-43x fewer bacterial reads with comparable execution time.

**Availability and implementation:** Hostile is implemented as an MIT licensed Python package available from https://github.com/bede/hostile together with supplementary material.

## 1 Introduction

Microbial specimens are often contaminated with host sequences. Since experimental host genome depletion protocols are imperfect, host DNA often reaches the sequencing instrument. Where the specimen host is a human, it is important that host sequences are subsequently deleted in order to protect anonymity. The widespread human contamination of publicly deposited microbial sequence data (Bush *et al*., 2020) is therefore regrettable and raises regulatory concerns, particularly in light of the rapid growth of metagenomic diagnostics. Furthermore, unwanted host sequences waste computing resources and may adversely affect downstream analyses such as variant calling and *de novo* assembly. Host decontamination is therefore the first step performed in many microbial genomic analyses. Existing approaches employ one of three strategies: *i)* exclusive retention of reads aligning to a target microbial genome (Hunt *et al*., 2022), *ii)* subtractive removal of reads aligning to a host genome, and *iii)* subtractive removal after metagenomic read classification (Kim *et al*., 2016; Wood et al., 2019). Where the target microbe is known *a priori*, the first strategy (exclusive retention) may be most suitable: for SARS-CoV-2 it is both more accurate and computationally efficient than subtractive removal (Hunt *et al*., 2022). However, the second and third strategies (subtractive removal) are generalisable, and thus necessary for analysis of microbes that are unknown *a priori*, mixtures, or novel.

In this article, we describe a simple tool implementing subtractive removal of contaminant human genome sequences, together with rigorous evaluation of its performance against real human genomes from the 1000 Genomes Project and simulated bacterial reads representing the 985 complete bacterial assemblies in the FDA-ARGOS dataset (Sichtig *et al*., 2019). We also report performance using simulated reads for 140 complete mycobacterial genomes. These results provide evidence of the accuracy of the approach in terms of both its ability to remove human host reads (sensitivity), and to retain microbial reads (specificity).

## 2 Materials and Methods

Hostile is implemented as a Python package providing a command line interface and Python API. The decontamination process involves a series of streaming operations on optionally gzip-compressed input FASTQ files: *i)* alignment to a custom human reference genome (Minimap2 or Bowtie2), *ii)* counting distinct reads (Samtools), *iii)* discarding aligned reads (and their mate reads for paired data; Samtools), *iv)* counting remaining reads (Samtools), *v)* Optionally replacing read names with incrementing integers (Awk), and *vi)* writing gzip-compressed FASTQ files (Samtools) (Li, 2018; Langmead and Salzberg, 2012; Danecek *et al*., 2021). These operations are streaming to reduce execution time and disk IO. Alignment of each read ceases after a single high quality match to the reference genome is found. For Bowtie2, default alignment parameters are used, while for Minimap2 a minimum chaining score of 40 is enforced for both long and short reads. Bowtie2 is the default aligner for short reads due to its relatively compact (<4GB) memory footprint, while Minimap2 is the default aligner for long reads, requiring approximately 12GB of random access memory (RAM) when using the map-ont preset for ONT (Oxford Nanopore Technologies) reads. Hostile generates summary statistics in JSON format including the total number of reads before and after decontamination. For ease of installation, Hostile is available as a Docker container and is packaged with Bioconda.

A custom human reference genome was built from the T2T-CHM13v2.0 human genome assembly (Nurk *et al*., 2022) and human leukocyte antigen (HLA) sequences. Human Illumina 2×100bp (Eberle *et al*., 2017) and ONT (Jain *et al*., 2018) reads from the well-characterised NA12878 sample were downsampled using BBTools (Bushnell, 2014) to a target depth of 10. We also examined 26 Illumina 2×150bp genomes at 30x depth representing each of the populations in the expanded 1000 Genomes Project, originating from Africa, Asia, Europe, and the Americas (Byrska-Bishop *et al*., 2022), in addition to newer data for the NA12878 sample. For FDA-ARGOS bacterial (*n*=985) and mycobacterial (*n*=140) metagenomes, Illumina reads were simulated with DWGSIM (Homer, 2010) while ONT reads were simulated with PBSIM2 (Ono *et al*., 2021). Refer to the Supplementary Text for detailed information about test data, masked reference genome construction, and read simulation.

We evaluated Hostile version 0.0.2 performance alongside the Human Read Removal Tool (HRRT; also known as Human Scrubber; https://github.com/ncbi/sra-human-scrubber) version 2.1.0. Testing was performed using an Ubuntu 22.04 AMD64 virtual machine. In order to address a defect in HRRT’s handling of paired reads and restore intended behaviour, BBTools was used to remove singleton reads from HRRT output.

## 3 Results

Full benchmark results are shown in Supplementary Tables S1 and S2, summarised in Table 1, and described here. Refer to the Supplementary Text for detailed information about test data preparation.

**Table 1.**
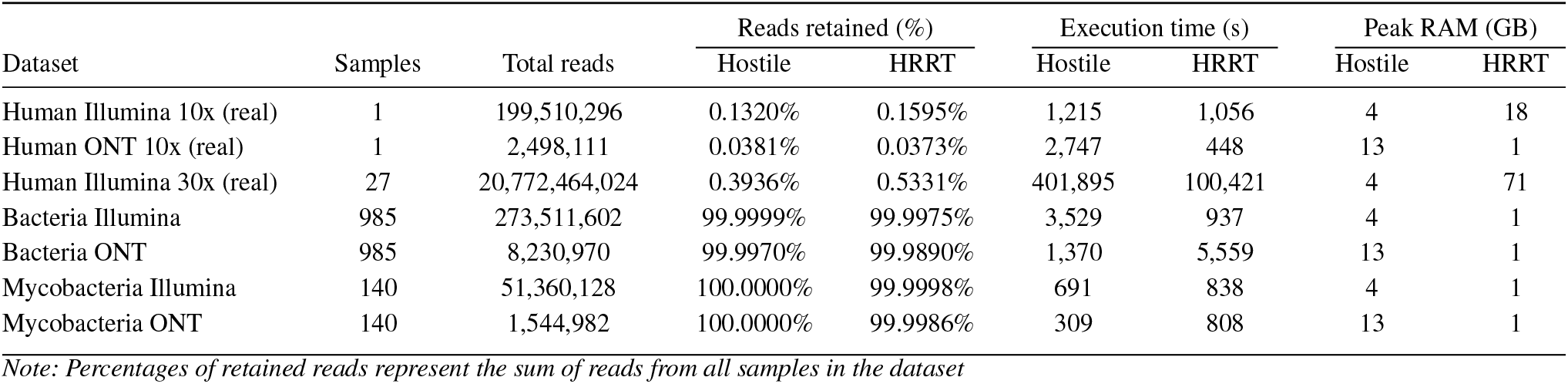
Evaluation of Hostile and the Human Read Removal Tool (HRRT) on real human and simulated bacterial and mycobacterial reads

### Accuracy of human read removal

For 10x depth 2×100bp Illumina data for NA12878, Hostile retained 0.132% of human reads while HRRT retained 0.160% (21% more). For 30x depth 2×150bp Illumina data for 27 representative genomes of each population in the expanded 1000 Genomes Projects (plus NA12878), Hostile retained 0.594% of human reads overall, while HRRT retained 0.729% (23% more). Surprisingly, human retention for NA12878 in these 30x 2×150bp data was considerably higher than in the older 10x 2×100bp data (0.132% vs. 0.641%). To investigate this discrepancy we performed *de novo* assembly (Li *et al*., 2015), revealing Epstein-Barr Virus (EBV) representation in all of the 27 studied Byrska-Bishop *et al*. genomes. Subtraction of EBV reads using Hostile with a custom index comprising accessions NC_007605.1 and NC_009334.1 enabled generation of adjusted accuracy figures, and highlighted one accession (ERR3242202) with 1.3% EBV contamination. Following EBV subtraction, retention for 2×150bp NA12878 decreased from 0.641% to 0.406% for Hostile and from 0.689% to 0.458%, with overall human retention for the 27 2×150bp genomes decreasing from 0.594% to 0.393%. For ONT data, Hostile retained 0.038% of reads while HRRT retained 0.037% (2% more). Using Hostile with a reference genome masked against bacterial sequences marginally increased retention of human reads from 0.131738% to 0.131979% (Illumina 2×100bp) and from 0.038029% to 0.038069% (ONT).

### Accuracy of microbial read retention

accuracy of bacterial read retention was evaluated using Illumina and ONT sequences simulated from reference-grade complete bacterial assemblies in the FDA-ARGOS dataset. For simulated Illumina data, Hostile retained 99.99989% of reads while HRRT retained 99.99752%, corresponding to HRRT removing 22 times as many bacterial reads as Hostile. Hostile’s bacterial read retention was further increased to 99.99994% through use of a reference genome masked against bacterial sequences. For simulated ONT data, Hostile and HRRT retained similar percentages of bacterial reads – 99.98918% and 99.98901% respectively. Use of a masked reference genome further reduced the number of bacterial ONT reads removed by Hostile from 891 to 251 (43 times less than HRRT). For mycobacterial reads, 140 complete assemblies were simulated in the same fashion. For simulated mycobacterial Illumina data, Hostile retained 99.99998% of reads while HRRT retained 99.999751%, corresponding to HRRT removing 15 times more mycobacterial reads than Hostile. For simulated mycobacterial ONT data, Hostile retained 99.99948% of reads while HRRT retained 99.99864%. Use of a masked reference with Hostile resulted in perfect (100%) retention of both Illumina and ONT reads.

### Execution time and memory usage

Execution time was measured as the median wall clock time required to process gzip-compressed FASTQ input and create gzip-compressed decontaminated FASTQ output with 8 threads. Three trials were performed for datasets other than the 27 2×150bp human Illumina genomes. See Table S1 for the commands used. Neither tool was faster for all datasets tested. HRRT was faster to decontaminate human reads, taking 1,056s and 448s to decontaminate real 2×100bp Illumina and ONT reads respectively, while Hostile took 1,215s and 2,743s. Decontaminating the 27 2×150bp human Illumina genomes with HRRT took 28hrs vs. 112hrs for Hostile. For simulated bacterial Illumina reads, HRRT was also faster, taking 937s vs. 3,507s for Hostile. However, for ONT reads, Hostile was faster, taking 1,369s vs. 5,559s for HRRT. For mycobacterial reads, Hostile decontaminated both Illumina and ONT reads faster than HRRT (687s and 309s for Hostile vs. 838s and 808s for HRRT). Memory (RAM) usage was measured with /usr/bin/time -v. Hostile consistently used 4GB when processing short reads (Bowtie2) and 13GB when processing long reads (Minimap2). In contrast, HRRT’s memory usage varied between 1GB and 71GB depending on the number of human reads present in the dataset, necessitating use of a virtual machine instance with 128GB of RAM in order to individually process the 27 2×150bp genomes from the 1000 Genomes Project.

## 4 Discussion

In any diagnostic or experiment where microbial genomes might be contaminated with human genomes, host decontamination is necessary both to safeguard patient anonymity and to avoid encumbering downstream analyses with redundant and potentially detrimental off-target sequences. For downstream analysis it is also of critical importance that microbial sequences are not inadvertently removed, leading to false negative variant calls and incomplete *de novo* assemblies. Where target microbes are unknown *a priori*, mixed or sufficiently novel, a subtractive human read removal approach is required, involving non-trivial computation using gigabytes of RAM. Hostile uses one of two complementary seed-and-extend aligners to accurately excise human reads. Bowtie2 is well-suited for decontaminating short reads due to its small memory footprint, fast index loading, and memory-mapped index support, while Minimap2 offers excellent long (and short) read performance in return for a larger index that is considerably slower to load. Compared to an existing approach, Hostile is more sensitive in terms of removing human reads, and an order of magnitude more specific in terms of retaining diverse bacterial reads, even without the use of a masked reference genome. Masked reference genomes can be easily created using a built-in utility, and prebuilt masked references are available to download.

While currently more accurate base-for-base, short reads present a greater challenge for decontamination due to their relatively low information content. Nevertheless, for a catalogue of short read genomes representing diverse human populations, Hostile removed 99.6% of reads. Although this figure accounts for widespread Epstein-Barr Virus contamination (>1% in ERR3242202), other non-human DNA likely accounts for a significant proportion of the remaining 0.4%. The figure of 99.6% should therefore be considered a lower bound for sensitivity with short reads.

Unlike existing tools, Hostile streams compressed FASTQ input to compressed FASTQ output without creating intermediate files. Hostile’s RAM requirements are increasingly met by consumer laptops, creating scope for accurate client-side host decontamination using what we hope will be broadly useful software.

## Data availability

International Nucleotide Sequence Database Collaboration (INSDC) accession numbers for sequencing datasets used in this article are provided in the supplementary material.

