## Supplementary Text for "Hostile: accurate host decontamination of microbial sequences"

Up to date information on installing and using this software can be found in the GitHub repository

#### Tables

1. Supplementary Table 1
2. Supplementary Table 2
3. Supplementary Table 3

#### Test data preparation

The 10x depth 2x100bp human datasets comprise reference quality real reads for NA12878 generated using Illumina HiSeq 2500 (ERR194147) and ONT (rel7) instruments respectively. These were downsampled to 10x target coverage using bbnorm.sh from BBTools:

```
bbnorm.sh target=10 in1=rel7.fastq.gz out=rel7.10x.fastq.gz
bbnorm.sh target=5 \
    in1=ERR194147.r1.fastq.gz \
    in2=ERR194147.r2.fastq.gz \
    out1=err194147_10x.r1.fastq.gz \
    out2=err194147_10x.r2.fastq.gz
```

The 30x depth 2x150bp human dataset of 27 samples comprises NA12878 and a single representative genome for each of the 26 populations characterised in the expanded 1000 Genomes Project. These are 2x150bp reads generated with the Illumina NovaSeq 6000. Accession numbers are provided in Supplementary Table 3

The datasets labelled 'Bacteria Illumina' and 'Bacteria ONT' comprise simulated reads at 10x depth for the 985 complete genomes in Database for Reference Grade Microbial Sequences as of 2023-06-01. Accession numbers are provided in Supplementary Table 3

The datasets labelled 'Mycobacteria Illumina' and 'Mycobacteria ONT' comprise simulated reads at 10x depth for 140 complete mycobacterial genomes whose accession numbers are provided in Supplementary Table 3

Illumina read pairs were simulated using dwgsim at 10x depth and 150bp in length with a random read probability (-y) of zero and somatic mutations (-F) disabled:

```
dwgsim -C 10 -1 150 -2 150 -y 0.0 -o 1 -z 1 -F 0.0 input.fasta
```

ONT reads were simulated using PBSim2 with a maximum length of 10000:

```
pbsim --depth 10 \
    --length-max 10000 --hmm_model P6C4.model input.fasta
```

The relevant software versions for test data preparation and benchmarking are as follows:

```
  - hostile=0.0.2
  - sra-human-scrubber=2.1.0
  - python=3.10.11
  - bedtools=2.31.0
  - biopython=1.81
  - pbsim2=2.0.1
  - minimap2=2.26
  - bowtie2=2.5.1
  - dwgsim=1.1.1
```

### Reference genome construction

A custom human reference genome was built by concatenating T2T-CHM13v2.0 and deduplicated IPD-IMGT/HLA v3.51.0 human leukocyte antigen sequences. This genome is automatically downloaded and cached to the user's application data directory (XDG_DATA_DIR) upon first execution of Hostile when using Hostile's Minimap2 backend. When Hostile's Bowtie2 backend is used, Hostile attempts to automatically retrieve a prebuilt Bowtie2 index constructed from this reference.

### Reference genome masking

150mers for each of the 985 complete FDA-ARGOS bacterial genomes and 140 complete mycobacterial genomes were created using a Python script accepting a reference sequence as an argument and generating FASTQ:

```python
import sys
from pathlib import Path
from Bio import SeqIO

def generate_fastq_kmers(ref_genome_file, read_length):
    with open(ref_genome_file, "r") as ref_fh:
        ref_stem = ref_genome_file.stem
        for record in SeqIO.parse(ref_fh, "fasta"):
            seq = record.seq
            for i in range(len(seq) - read_length + 1):
                read = seq[i:i+read_length]
                print(
                    f"@{ref_stem}_pos_{i}\n"
                    f"{read}\n"
                    f"+\n"
                    f"{'?'*read_length}"
                )
generate_fastq_kmers(Path(sys.argv[1]), read_length=150)
```

FASTQ files of the 150mers for each bacterial genome were concatenated and aligned to a human reference genome, converted to BED coordinates which were then applied to the unmasked genome, creating the masked genome.

```
time minimap2 -ax map-ont human.fa 150mers.fastq.gz \
    | samtools sort - > 150mers.bam
bedtools bamtobed -i 150mers.bam > 150mers.bed
bedtools maskfasta -fi human.fa -bed 150mers.bed -fo human-masked.fa
```

As of version 0.0.3, the command line interface includes a mask subcommand for easily and quickly masking arbitrary reference genomes with arbitrary masking targets without the need for simulation.
